# Reactivation of early-life stress-sensitive neuronal ensembles contributes to lifelong stress hypersensitivity

**DOI:** 10.1101/2022.12.21.521303

**Authors:** Julie-Anne Balouek, Christabel Mclain, Adelaide R. Minerva, Rebekah L. Rashford, Shannon N. Bennett, Catherine Jensen Peña

**Affiliations:** Princeton Neuroscience Institute, Princeton University, Princeton, NJ 08544, USA

## Abstract

Early-life stress (ELS) is one of the strongest lifetime risk factors for depression, anxiety, suicide, and other psychiatric disorders, particularly after facing additional stressful events later in life. Human and animal studies demonstrate that ELS sensitizes individuals to subsequent stress. However, the neurobiological basis of such stress sensitization remains largely unexplored. We hypothesized that ELS-induced stress sensitization would be detectable at the level of neuronal ensembles, such that cells activated by ELS would be more reactive to adult stress. To test this, we leveraged transgenic mice to genetically tag, track, and manipulate experience-activated neurons. We found that in both male and female mice, ELS-activated neurons within the nucleus accumbens (NAc), and to a lesser-extent the medial prefrontal cortex, were preferentially reactivated by adult stress. To test whether reactivation of ELS-activated ensembles in the NAc contributes to stress hypersensitivity, we expressed hM4Dis receptor in control or ELS-activated neurons of pups and chemogenetically inhibited their activity during experience of adult stress. Inhibition of ELS-activated NAc neurons, but not control-tagged neurons, ameliorated social avoidance behavior following chronic social defeat stress in males. These data provide evidence that ELS-induced stress hypersensitivity is encoded at the level of corticolimbic neuronal ensembles.

## INTRODUCTION

Experience of early-life stress (ELS) is a major risk factor for mood and anxiety disorders later in life, with an effect size that far outweighs any known genetic risk (Green et al., 2010; Scott et al., 2012). Each year, nearly 700,000 children in the United States experience child maltreatment such as neglect or abuse (Stedt, 2018). Many more children experience other forms of profound stress including death of a caregiver, parental incarceration, displacement due to natural disasters or violence, and socioeconomic disadvantage, indicating a genuine need for a better understanding of the long-lasting neurobiological consequences of childhood maltreatment and stress (Hackman et al., 2010). Studies in humans and animal models indicate that ELS increases risk for anxiety and depression by sensitizing individuals to stress later in life, leading to a first appearance or synergistic worsening of depression-like symptoms after additional stress (McGuinn et al., 2005; McLaughlin et al., 2010; Saxton & Chyu, 2020; Sidamon-Eristoff et al., 2022; Zhang et al., 2016). The brain is particularly sensitive to environmental experiences when it is rapidly developing early in life, and evidence suggests that there may be sensitive windows in postnatal development when childhood adversity increases vulnerability to depression (Dunn et al., 2018, 2019; Peña et al., 2017, 2019). However, the cellular mechanisms linking early adverse experience with enhanced sensitivity to stress and vulnerability to psychiatric disease remain largely unknown.

Experience has been shown to facilitate formation or strengthening of cellular ensembles which are reactivated during recall of the experience (Cai et al., 2016; Deng et al., 2013; Han et al., 2009; Josselyn et al., 2015; Kitamura et al., 2017; Liu et al., 2012; Pignatelli et al., 2019; Ramirez et al., 2013; Ryan et al., 2015; Tonegawa, Liu, et al., 2015; Tonegawa, Pignatelli, et al., 2015). In addition to the pioneering findings in the hippocampus (Guzowski et al., 1999; McKenzie et al., 2014), subsequent studies have also shown the sufficiency, necessity, and correlation of putative neuronal ensembles for a fear memory in several other brain structures including regions of the amygdala, medial prefrontal cortex, hypothalamus, and retrosplenial cortex, demonstrating that cell ensembles for a given memory are not restricted to a single brain region but may be dispersed extensively throughout the brain (Butler et al., 2015; Cowansage et al., 2014; Davis & Reijmers, 2018; Han et al., 2007; Janak & Tye, 2015; Matos et al., 2019; Roy et al., 2022; Ryan et al., 2015). Moreover, there is evidence that distinct types of memory, such as fear, reward, and social memories, are encoded in distinct ensembles within the brain (Hsiang et al., 2014; McKenzie et al., 2014; Okuyama et al., 2016; Redondo et al., 2014; Shpokayte et al., 2022; Zhou et al., 2019). Reactivation of ensembles associated with either positive or negative valence experiences is sufficient to drive preference or aversion in a reversible manner (Shpokayte et al., 2022), and ablating or inhibiting fear-associated ensembles of cells erases their associated fear memories (Denny et al., 2014; Han et al., 2009). We hypothesized that one mechanism of ELS-induced stress hypersensitivity was through reactivation of ELS-sensitive cellular ensembles in mesocorticolimbic regions of the brain.

While there are many tools to record neuronal activity in real-time, until the last decade identifying and accessing cells activated by a distant past experience has been a major challenge in the field of neuroscience (DeNardo & Luo, 2017). To overcome this challenge, we took advantage of advances in mouse transgenics that allow us to genetically tag, track, and manipulate experience-activated cells in a temporally-specific manner. Using transgenic mice (*Arc-CreER^T2^*) to indelibly label ELS-activated cells (Denny et al., 2014), we investigated the recruitment and reactivation of ELS-activated cellular ensembles during adult stress as indicated by fluorescent imaging of single- and double-labeled cells throughout key mesocorticolimbic regions known to be sensitive to stress across the lifespan (Russo & Nestler, 2013; Hanson et al., 2021). We specifically focused on the nucleus accumbens (NAc) and its neuromodulatory and glutamatergic inputs including ventral tegmental area (VTA), basolateral amygdala (BLA), and medial prefrontal cortex (mPFC). This approach allowed us to test two possible hypotheses: 1) that adult stress activates overall more neurons given a history of prior ELS; and 2) that ELS-activated cells are preferentially re-activated by adult stress. Finally, using Cre-dependent designer receptors exclusively activated by designer drugs (DREADD) chemogenetic manipulation within these mice, we silenced ELS-activated ensembles in the NAc during adult stress to test the hypothesis that cells initially activated by ELS contribute to behavioral hypersensitivity to future stress.

## METHODS

### Mice

All protocols were performed in compliance with the Guide for the Care and Use of Laboratory Animals (NIH, Publication 865–23) and were approved by the Institutional Animal Care and Use Committee of Princeton University (IACUC protocol #2135). Mice were housed in a temperature- and humidity-controlled animal care facility with 12 h light/ dark cycle (lights on at 12:00 am) and group-housed in sets of 3-5 with *ad libitum* access to food and water. Mice were considered to be “male” or “female” based on external genitalia (adult) and/or anogenital distance (pups). For all experiments, mice were bred in-house in trios. Males were removed after 5 days, and females remained housed together until 19 days post-mating, at which point females were individually housed in clean cages with nesting material from the previous cage for olfactory continuity to reduce stress. Litters were randomly assigned at birth to standard-rearing (Std) or early life stress (ELS) conditions, described below. All offspring were weaned at postnatal day P21 by sex. Both males and females were included in all experiments.

Transgenic mice were on a C57BL/6J background. Heterozygous *Arc-CreER^T2/+^* mice (Denny et al., 2014); JAX stock #022357) were bred with *R26-CAG-LSL-Sun1-sfGFP-Myc* knockin mice (“Sun1-sfGFP;” JAX stock #030952; Mo, et al. 2015) for imaging experiments. In this cross, recombination and expression of the *Sun1-sfGFP* transgene is activity- and ligand-dependent. Activity-induced expression of the immediate early gene Arc drives expression of the Cre-ERT2 fusion protein, which is sequestered in the cytoplasm until injection of a selective estrogen receptor modulator such as 4-hydroxytamoxifen (4-OHT). Administration of 4-OHT allows relocalization of CRE to the nucleus, where it can act as a recombinase to remove the floxed STOP cassette and allow permanent expression of the fusion SUN1-sfGFP protein. SUN1 is an inner-nuclear membrane spanning protein, such that SUN1-sfGFP becomes localized to the nuclear membrane of neurons activated during 4-OHT’s duration of action. Double-transgenic offspring were used for all imaging studies. Offspring of heterozygous *Arc-CreER^T2/+^* bred in-house with C57Bl/6J wild-type mice were used for DREADD experiments. Wild-type C57Bl/6J mice were used for DREADD ligand dose piloting studies. Retired Swiss-Webster male breeder mice (Taconic) were housed individually and used as aggressor mice in chronic non-discriminatory social defeat stress.

### Genotyping

Offspring genotyping was done by toe clip within 1 day of birth. *Arc-CreER^T2/+^* × *Sun1-sfGFP* founders and subsequent offspring were genotyped using the following primer sets: *Cre* 5’-GCC TGC ATT ACC GGT CGA TGC AAC G-3’; 5’-AAA TCC ATC GCT CGA CCA GTT TAG TTA CCC-3’; *Sun1-sfGFP* (mutant): 5’-CTG AAC TTG TGG CCG TTT AC-3’; 5’-ACA CTT GCC TCT ACC GGT TC-3’; (wild-type) 5’-CAG GAC AAC GCC CAC ACA-3’; and 5’-AAG GGA GCT GCA GTG GAG TA-3’. *Cre* and *Sun1-sfGFP* genotypings were performed separately. *Sun1-sfGFP* genotyping was performed following The Jackson Laboratory genotyping protocol.

### Early life stress paradigm

ELS occurred from P10-17 as previously described (Peña et al., 2017, 2019) and consisted of both maternal separations (with all pups from a litter removed to a clean cage and returned to the home cage 3-4 hours later) and limited nesting material in the home cage (to 1/3 standard EnviroDri nesting puck size). Pups were given access to diet gel and several chow pellets on the cage floor during separations, although pups were not observed to eat anything. At the conclusion of ELS on P17, mice were given clean cages with a full puck of nesting material, and left undisturbed until weaning. Standard facility-reared pups (Std) were reared with a normal amount of nesting material and left unhandled.

### Administration of 4-hydroxy-tamoxifen (4-OHT)

(Z)-4-Hydroxytamoxifen (4-OHT; Sigma-Aldrich #H7904) was first prepared as 25 mg/mL stock solutions in ethanol and frozen at −20C. For injections, 4-OHT was prepared fresh daily from stock to 5 mg/mL, dissolved in ethanol and corn oil (1:4). At P17, double-transgenic *Arc-CreER^T2/+^* × *Sun1-GFP* mice received intraperitoneal injections at 50 mg/kg body weight. Injections were given to ELS pups at the start of four hours of maternal separation. In order to tag a specific, positive valence experience in standard-reared pups, pups were provided with a novel enrichment object (a plastic exercise saucer) in the home cage at P17 prior to 4-OHT administration. Pups were observed to explore the novel object but not necessarily run on it.

### Adult Stress: Chronic Non-discriminatory Social Defeat Stress (CNSDS)

Both male and female mice were exposed to chronic non-discriminatory social defeat stress (CNSDS) (Dieterich et al., 2021; Yohn et al., 2019), a modified version of chronic social defeat stress (Berton et al., 2006) for simultaneous social stress of male and female mice. Briefly, beginning at age P60, CNSDS-assigned mice were subject to 10 days of daily social defeat by a Swiss Webster retired breeder (“aggressor;” Taconic) in a standard rat cage filled with corn cob bedding. An experimental male mouse was introduced to an aggressor’s cage first for ~3 minutes, followed by an experimental female mouse for an additional 5 minutes. Males were then moved across a perforated plexiglass barrier within the aggressor cage, while females were single-housed with a handful of soiled aggressor bedding (imaging experiments) or co-housed in a new cage across a plexiglass barrier from a different aggressor (DREADD experiments). Experimental mice were introduced to a new aggressor each day for 10 days, and males and females rotated in different directions so that trios were unique each day. Control mice were housed in a standard mouse cage in pairs of the same sex, separated by a perforated plexiglass barrier. For imaging experiments, mice were sacrificed one hour after the last bout of social defeat.

Mice in the DREADD experiment underwent CNSDS twice (**Figure 5C**). In the first round, all mice received a vehicle injection of normal saline (i.p.) 30-minutes prior to social defeat or handling (control mice). In the second round, all mice received ligand injection (see below) 30-minutes prior to social defeat or handling. Notably, for DREADD experiments, both male and female experimental mice were only co-housed with aggressors for a maximum of 4 hours after each day of social defeat to account for the 3-5-hour functional period of OLZ (Aravagiri et al., 1999; Mattiuz et al., 1997) so that presence of the aggressor did not continue to stress mice after DREADD-inhibition of neurons was expected to wear off. At the conclusion of each round of CNSDS, male and female mice were individually housed in clean cages for the duration of behavior testing.

### Immunohistochemistry and imaging

Mice were sacrificed one hour after the last bout of early-life stress (Figure 2) or adult stress (Figures 3–4) to capture maximal stress-induced cFos levels. An additional four adult *Arc-CreER^T2/+^* × *Sun1-sfGFP* mice which never received 4-hydroxytamoxifen were sacrificed directly from the home cage in order to assess background “leak” of Sun1-GFP expression (Figure 1). Mice were anesthetized with ketamine/xylazine (100 mg/kg and 10 mg/kg, i.p.) and transcardially perfused with sterile 1X PBS followed by 4% paraformaldehyde. Brains were removed, equilibrated in 30% sucrose in PBS, and frozen at −80°C until processing. Brains were sectioned at 50 μm-thick slices using a cryostat. Immunohistochemistry was performed using the following antibodies: cFos (Cell Signaling, clone9F6, mAb#2250), GFP (Aves Lab, GFP-1020), tyrosine hydroxylase (ImmunoStar, 22941), mcherry (DREADD) (Abcam, ab167453) and Prolong Gold Antifade with DAPI (Invitrogen). Imaging was performed on Hamamatsu NDP slide scanner (Hamamatsu Nanozoomer 2.0HT). Detection, colocalization, and quantification were performed using ndpisplit (Deroulers et al., 2013) and ComDet v.0.5.3 plugin for FiJi/ImageJ (https://github.com/ekatrukha/ComDet) with the following parameters: Pixel size=12, Intensity threshold=9 for both channels. Imaging data: each value is a mean of 6 traced ROIs for each animal on average (minimum of 2, maximum of 8), such that sample size is based on subjects rather than slices (*n*=2-4 mice/group for validation imaging; 4-11 mice/ group for reactivation imaging).

**Figure 1.**
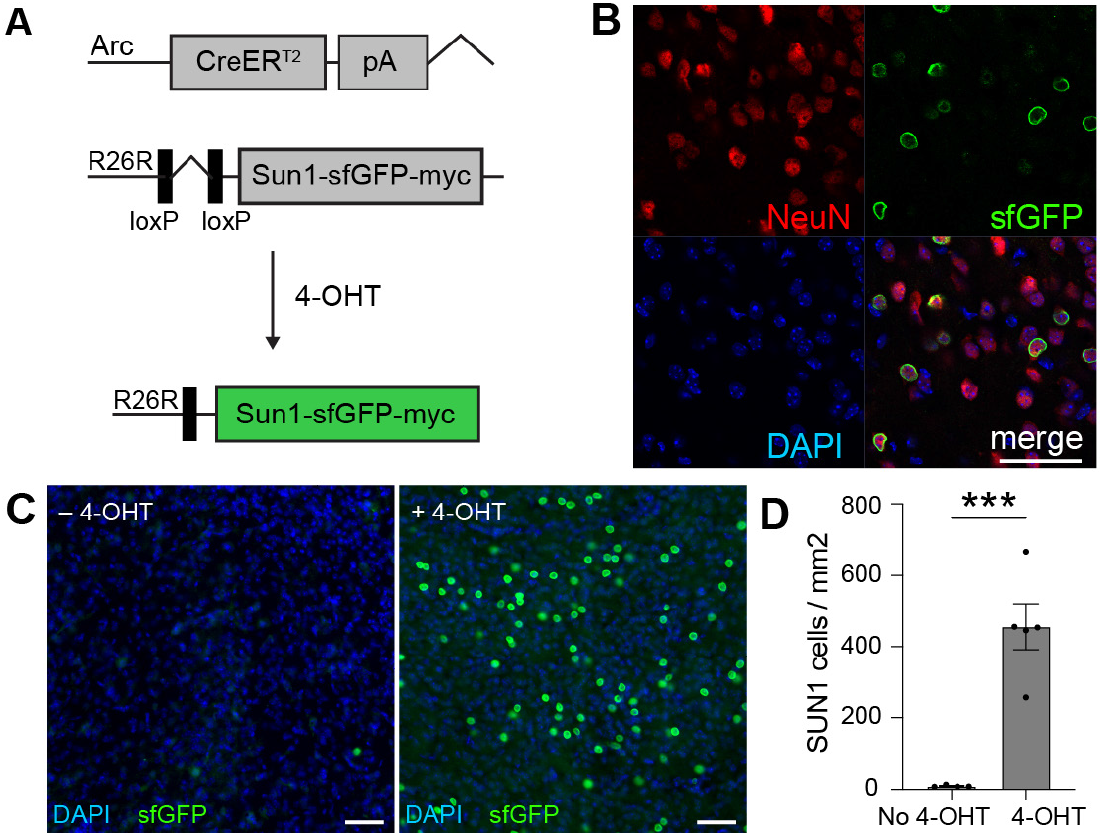
Validation of activity-dependent transgenic mice show neuronal specificity and low leak. (A) Schematic for 4-OHT-dependent and Arc-expression-dependent recombination and transgene expression in double-transgenic *Arc-CreER^T2/+^* × *Sun1-sfGFP* mice. (B) Representative 63x image within NAc of *Arc-CreER^T2/+^* × *Sun1-sfGFP* mice after recombination, labeling all cells (DAPI), all neurons (NeuN), and previously-activated cells (GFP). The scale bar represents 50 μm. (C) Representative 20× images within NAc of *Arc-CreER^T2/+^* × *Sun1-sfGFP* adult mice without (left) or with (right) prior 4-OHT administration at P17 show that spontaneous recombination without 4-OHT (“leak”) is low. The scale bar represents 50 μm. (D) Quantification of GFP+ cell density in double-transgenic mice with or without prior 4-OHT-induced recombination.

**Figure 2.**
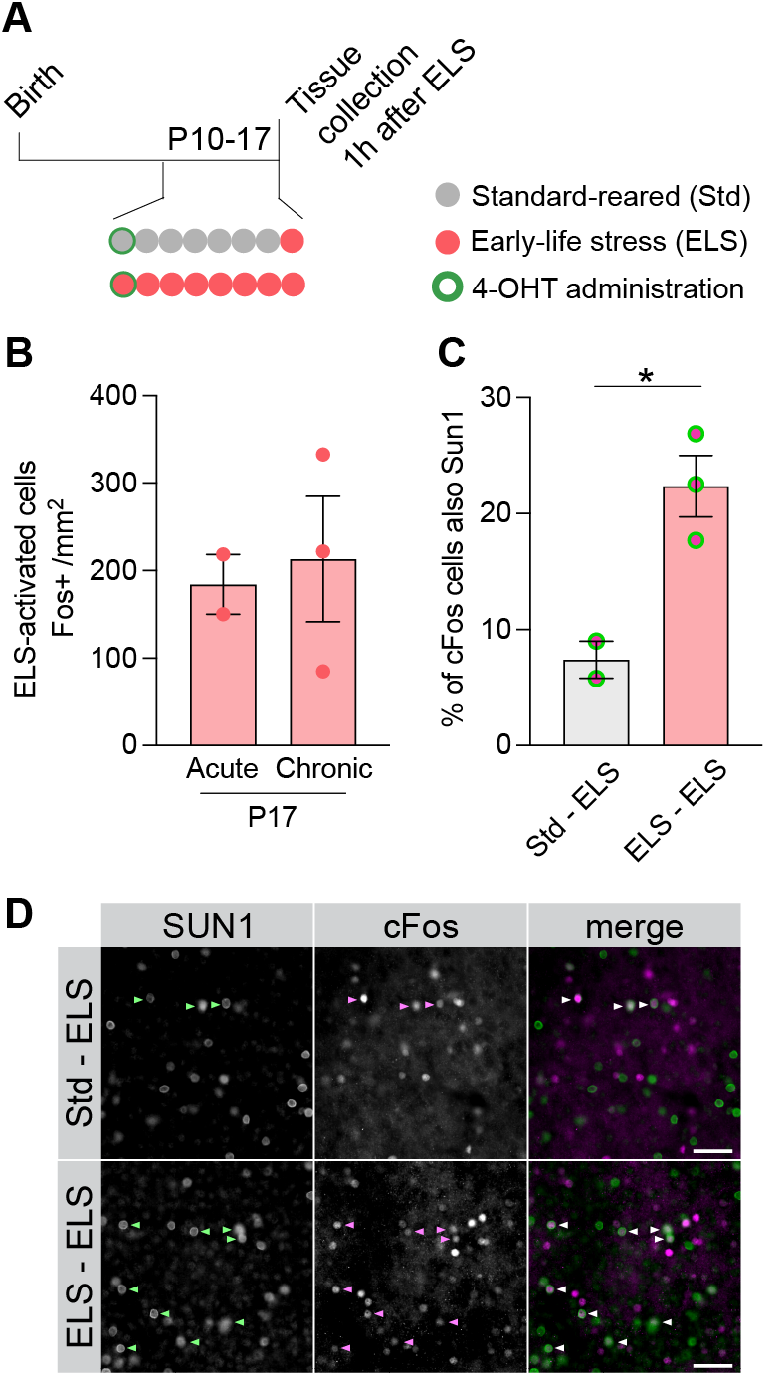
Experience-dependent neuronal labeling is specific. (A) Experiment design for assessing specificity and drift of ELS activity: genetic recombination was induced with 4-OHT at P10 and tissue was collected on P17. “Acute” stress mice were presented with enrichment on the day of tagging at P10, were reared in standard conditions, and then exposed to a single day of ELS on P17. “Chronic” stress mice were subjected to ELS daily from P10 to P17. (B) No changes were observed in the density of ELS-activated cells in NAc following either acute or chronic stress, as assessed by cFos+ cell density in response to ELS at P17. (C) Quantification of the percent of cFos+ cells active in NAc during ELS at P17 that were also active (GFP+) during either Std/enrichment or ELS at P10 revealed significantly greater reactivation under matched conditions (ELS-ELS) compared to unmatched conditions (enrichment vs. ELS). (D) Representative images in NAc associated with C. The scale bar represents 50 μm.

**Figure 3.**
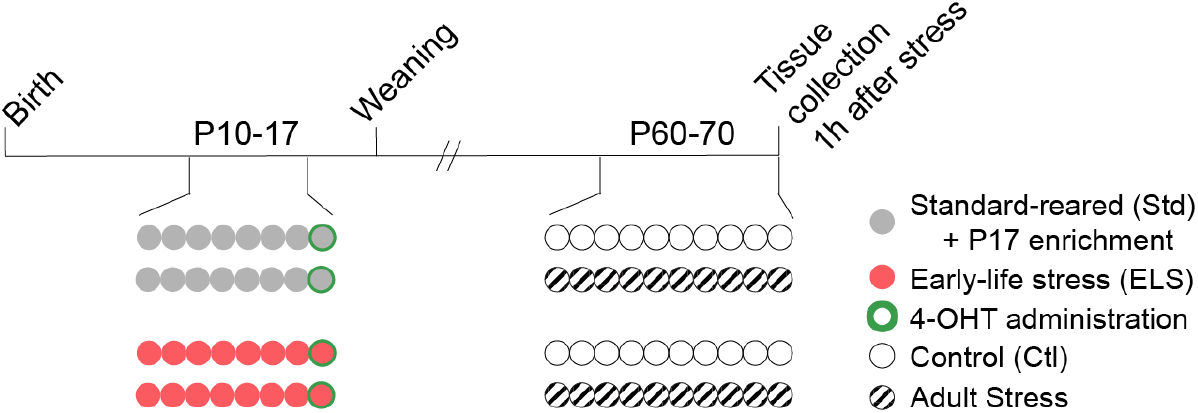
Experimental design for activity-dependent ELS-engram labeling. *Arc-CreER^T2/+^* × *Sun1-sfGFP* mice were either Std-reared or experienced ELS from P10-17. Recombination was induced by 4-OHT administration on P17 prior to experience with a novel exercise saucer in the home-cage (Std mice) or maternal separation (ELS). In adulthood, half of each group were assigned to 10 days of control conditions or CNSDS, and tissue was collected one hour after the final bout of social defeat stress.

**Figure 4.**
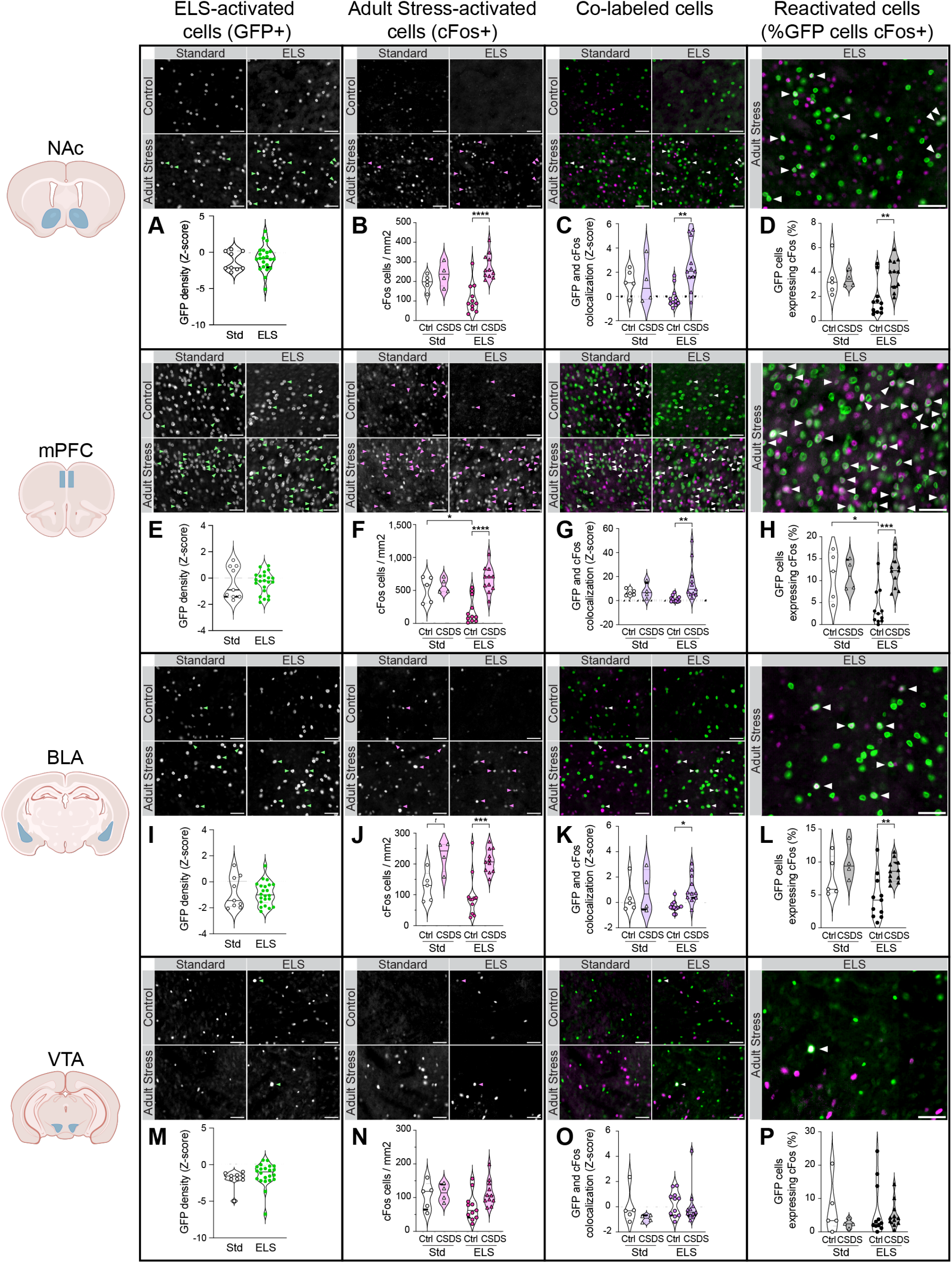
Stress-specific neuronal activation and reactivation in mesocoticolimibic brain regions. Each panel contains representative 20× images (scale bar = 50 μm) of mice from four groups (Std-Ctrl, ELS-Ctrl, Std-CNSDS, or ELS-CNSDS), and their quantification. Arrows indicate co-localized GFP+ and cFos+ cells. Four brain regions of the reward pathway were assessed: (A-D) are measures in NAc, (E-H) in mPFC, (I-L) in BLA and (M-P) in VTA. (A), (E), (I), and (M) show Std or ELS-activated cell density (GFP+), presented as a Z-score to correct for increased labeling in Sun1/Sun1 mice compared to Sun1/+ mice. (B), (F), (J), and (N) show control or CNSDS-activated cell density (cFos+). (C), (G), (K), and (O) show a Z-score of the density of co-labeled cells (GFP+ and cFos+). (D), (H), (L), and (P) show the percentage of Std or ELS-activated cells that were reactivated during adult stress [100*(GFP+ and cFos+)/GFP+].

### Behavioral testing

A within-subject design was employed to test adult male and female behavior three times: first before CNSDS, after a first round of CNSDS with vehicle administration, and after a second round of CNSDS with DREADD ligand (**Figure 5C**).

**Figure 5.**
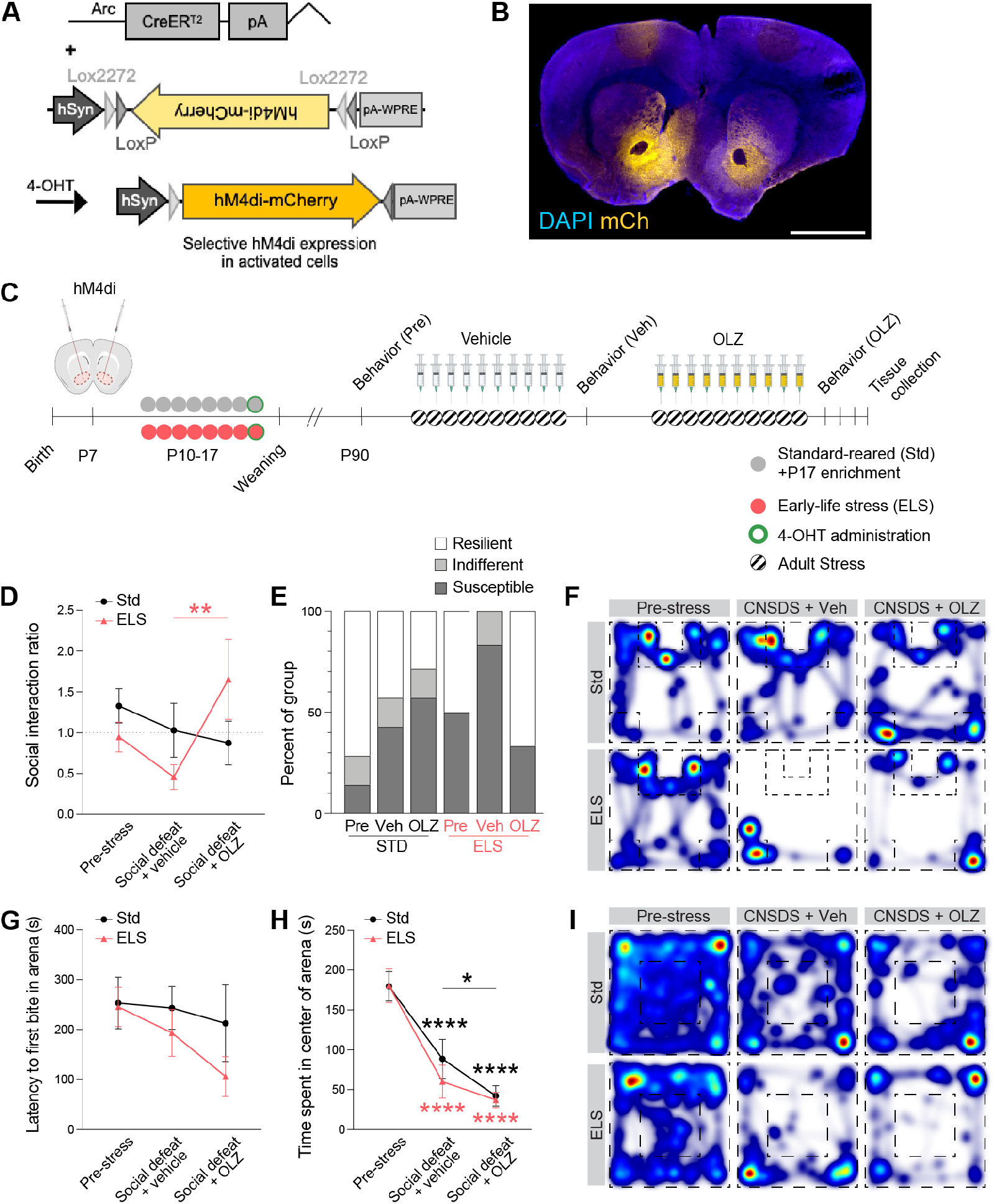
Inhibition of early-life stress-activated cells in NAc reduces hypersensitivity to adult stress. (A) Viral expression strategy: Administration of 4-OHT to *Arc-CreER^T2/+^* mice injected with AAV-hSyn-hM4Di-mCherry results in permanent hM4Di expression in experience-activated cells. (B) Representative image of DREADD-mCherry expression in NAc from pups infused with virus on P7 shows relative specificity of injection even within pups. The scale bar represents 2 mm. (C) Timeline and experimental groups: Mice were either Std-reared or subjected to ELS from P10 to P17. Std-reared mice were given access to a running wheel before 4-OHT-induced recombination on P17. In adulthood, all mice underwent two rounds of CNSDS for 10 days. During the first round, all mice received a vehicle injection 30 minutes prior to each daily bout of social defeat; in the second round all mice received 0.1 mg/kg OLZ 30 minutes prior to social defeat. Mice were assessed behaviorally before and after each round of CNSDS. (D) Social interaction ratio among male mice, where lower ratio indicates greater social avoidance. (E) Percent of male mice categorized as resilient, indifferent, or susceptible to CNSDS under each treatment condition. (F) Representative heat maps of behavior in each group during social interaction testing with aggressor present. (G) Latency to eat in a novel arena during a novelty-suppressed feeding test. (H) Time spent in the center during open field testing. (I) Representative heat maps of behavior in each group during open field exploration. OLZ indicates DREADD-inhibition of early experience-activated cells. Post-hoc comparisons between corresponding conditions and pre-stress data are indicated by **(*p*<0.01) and ****(*p*<0.0001).

### Social interaction

To assess social avoidance behavior, experimental mice were tested in a two-stage social interaction test under red lighting, as previously described (Peña et al., 2017). Social avoidance has been previously associated with other depression-like behaviors and is responsive to antidepressant treatment (Berton et al., 2006; Krishnan et al., 2007). In the first 2.5-min stage, the experimental mouse was allowed to freely explore an arena (44 × 44 × 20 cm) containing a novel plexiglass and wire mesh enclosure (novel object; 10×6 cm) centered against one wall of the arena. In the second 2.5 min stage, the experimental mouse was immediately returned to the arena with a novel Swiss Webster mouse (aggressor strain) within the enclosure. Time spent in the ‘interaction zone’ (14×26 cm) surrounding the enclosure, ‘corner zones’ (10×10 cm), and ‘distance traveled’ within the arena was measured by video tracking software (Ethovision, Noldus). A social interaction ratio (SI Ratio) was calculated of time spent exploring the novel mouse over time exploring the novel object; mice were considered “susceptible” to CNSDS if SI Ratio<0.9, “resilient” if SI Ratio >1.1, and “indifferent” for interaction scores in between (Peña et al., 2017). To combat habituation during repeated testing, patterns on the walls of the chambers were altered for each of the three testing sessions.

### Open-field test

Mice were allowed to explore a brightly lit, empty arena (44 × 44 × 20 cm) for 10 minutes. Total distance traveled, velocity and time spent in the center (20 × 20 cm) were recorded and measured via Ethovision (Noldus), as previously described (Peña et al., 2017). To combat habituation during repeated testing, patterns on the walls of the chambers were altered for each testing session.

### Novelty-suppressed feeding

Novelty suppressed feeding testing was adapted from previous work (Bodnoff et al., 1989). Mice were food deprived for 18 hours in a clean homecage. On testing day, they were placed in a corner of a novel, brightly-lit (200 lux) arena (44 × 44 × 20 cm). The floor was covered with corncob bedding and the arena walls were covered with a geometric pattern to enhance the novelty of the environment. A single food pellet was attached to a white circular platform (10 cm diameter) in the center of the arena. The test lasted until mice took their first bite (sitting on their haunches and using their forpaws to hold and bite the pellet), or for a maximum of 10 minutes. Mice that timed out were assigned a latency of 600 seconds. Immediately after taking their first bite, mice were transferred to their home cages and given access to their usual amount of food. To combat habituation during repeated testing, patterns on the walls of the chambers were altered for each testing session.

### Pharmacogenetic inhibition of cellular activity

Designer receptors exclusively activated by designer drugs (DREADD) were used to suppress cellular activity. Viral AAV transgene expressing Cre-dependent inhibitory DREADD (pAAV-hSyn-DIO-hM4D(Gi)-mCherry) was injected into NAc by aseptic intracranial stereotaxic surgery in *Arc-CreER^T2/+^* pups at postnatal day P7 [Addgene #44362; originally gifted from Bryan Roth; http://n2t.net/addgene:44362 (Krashes et al., 2011)]. All mice received hM4D(Gi) transgenes and thus controls were within-mouse based on vehicle/ligand administration.

At P7, *Arc-CreER^T2/+^* pups were anesthetized with isoflurane (2-3% induction; 1-2% maintenance) for surgeries and fitted into dual-arm stereotax with the nose fitted in the nose cone above the bite bar, and the head stabilized using blunt ends of ear bars between the ear and jaw. To bilaterally target the nucleus accumbens (NAc), two Hamilton syringes were angled inward 7° and the following coordinates were used: ML: ± 1.4 mm, AP: 1.1 mm, DV: −4.2 mm. 200 nL of AAV (2.5 × 1013) were injected on each side. Both VetBond glue and two sterile sutures were used to ensure complete healing of incisions. Mice were given a perioperative topical dose of bupivacaine (0.25%, 2 mg/kg) and an additional dose 24h following surgery. To prevent cannibalization of pups by dams following surgery, pups recovered in partially-warmed cage and dams were habituated with a piece of paper towel with sterile eye lubricant (Puralube Vet Ointment Sterile Ocular Lubricant; Dechra) and liquid tissue adhesive (VetBond, 3M); pups were only returned to the home cage with the dam once completely ambulatory. At P17, after 10 days of transgene expression, recombination was induced with 4-OHT as described above.

Olanzapine (OLZ; Henry Schein) was chosen as the hM4Di ligand for inhibition of cellular activity as it has been shown to cross the blood-brain barrier, unlike the more common DREADD ligand clozapine N-oxide (CNO), and to be a specific and potent activator of hM4Di (Weston et al., 2019; Upright and Baxter, 2020). We sought to avoid CNO as it is back-converted to clozapine, which has unfavorable side-effects in humans and may impact anxiety-like behavior at effective doses in mice. A dose of 0.1 mg/kg body weight OLZ was previously reported to be an effective ligand for hM4di DREADDs *in vivo* in mice (Weston et al., 2019). OLZ was prepared fresh daily from a −20C stock and diluted in normal saline to working concentrations (5 mg/mL). First, we tested two doses of OLZ (either 0.1 or 1.0 mg/kg body weight, i.p., given 30-minutes prior to behavioral testing) for a potential impact on anxiety- and depression-like behavior and weight gain in wild-type C57Bl6/J mice without hM4di expression (*n*=6 male mice/group). A dose of 0.1 mg/kg body weight was selected for subsequent experiments based on lack of impact on behaviors of interest in the absence of DREADD. For DREADD inhibition experiments, both control and CNSDS mice were administered vehicle (sterile normal saline) or 0.1 mg/kg OLZ (i.p.) 30-minutes prior to handling or social defeat, respectively, for the duration of CNSDS (10 days) (*n*=8-9 male and female mice/group).

### Data analysis and statistics

All data are plotted at Mean ± the standard error of the mean (SEM). We observed a small but significant effect of Sun1/+ vs. Sun1/Sun1 genotype, with more cells counted in homozygous mice; to correct for this, Z-scores of GFP+ cells were calculated for Sun1/+ and Sun1/Sun1 genotypes separately. GFP and colocalization quantifications are thus presented as z-scores compared to the standard group. For imaging data, counts from 2-6 slices per subject were averaged and the mean per subject was plotted and used for analysis, such that *n* represents individual subjects rather than replicate slices. All statistics were performed using SPSS (Version 26) or Prism (GraphPad; version 9), with alpha set to 0.05. Student’s t-test was used to compare two groups. Two-way ANOVA was used to test main effects and interactions of early-life and adult stress. Behavioral effects were assessed using repeated-measures ANOVA. Post-hoc analyses between pairs of groups were conducted using Tukey’s multiple-comparisons correction if there were significant main effects or interactions. Significant changes in proportion of resilient mice in the social avoidance task were calculated by two-tailed two-proportion z-test.

### Data Availability

All relevant data that support the findings of this study are available by request from the corresponding author (CJP).

## RESULTS

### Validation of activity-dependent transgenic mice

To validate whether our double-transgenic *Arc-CreER^T2/+^* × *Sun1-sfGFP* mice (**Figure 1A**) allowed for an activity-dependent, ligand-dependent label with minimal background expression, we examined transgene expression with and without administration of 4-OHT in a homecage context. Administration of 4-OHT to Std-reared mice in adulthood produced sfGFP localized to the nuclear membrane within one week. Consistent with known neuronal expression of Arc (Pastuzyn et al., 2018), all sfGFP-expressing cells colocalized with NeuN staining, confirming neuronal specificity (**Figure 1B**). Absence of 4-OHT administration produced very little sfGFP expression even in adulthood, indicating low levels of spontaneous genetic recombination and “leakage” [*t*(1,7)=6.103, *p*=0.0005, **Figures 1C-D**].

### Experience-dependent neuronal labeling is specific

Previous work has shown that rodents mount a similar corticosterone response to maternal separation across days of separation, indicating a lack of habituation, but the extent to which mesocorticolimbic neuron activity might habituate to repeated ELS was not yet known. To test this, and the specificity of activity labeling to experience, we administered 4-OHT to *Arc-CreER^T2/+^* × *Sun1-sfGFP* pups on P10 to label neurons during ELS or a distinct experience (home-cage enrichment for 4 hours in Std-reared mice). Pups continued to experience either ELS or Std conditions, and then all mice experienced ELS on P17 one-hour prior to tissue collection (**Figure 2A**). Quantification of cFos+ cells in NAc on P17 one hour after experiencing ELS either for the first time (“acute”) or eighth time (“chronic”) revealed similar levels of active neurons (**Figure 2B**), indicating that mice do not habituate to ELS experience at the level of NAc activity.

We next sought to understand whether ELS activated the same ensembles of neurons across days, and the specificity of such activity. Quantification of the overlap of cFos+ neurons activated by ELS on P17 that were previously active during P10 enrichment or P10 ELS revealed significantly greater reactivation under matched conditions [*t*(1,3)=4.122, *p*=0.026; **Figures 2C-D**], suggesting relative specificity and stability of ELS-sensitive ensembles in NAc, even after one week.

### Mesocorticolimbic neurons activated by early life stress, adult stress, or both

We next sought to investigate two possible hypotheses for how ELS-induced hypersensitivity to adult stress may be encoded at a cellular level: 1) that adult stress activates overall more neurons given a history of prior ELS; and 2) that ELS-activated cells are preferentially re-activated by adult stress. To test these hypotheses, we used *Arc-CreER^T2/+^* × *Sun1-sfGFP* mice and a 2×2 experimental design of ELS and/or adult chronic non-discriminatory social defeat stress (CNSDS). With this design, Arc-driven SUN1-sfGFP expression represented cells active during specific early life experience and expression of the immediate early gene cFos at sacrifice represented cells active during adult experience (**Figure 3**). In order to tag a specific, positive valence experience in Std-reared pups, and to tag a similar number of neurons in Std and ELS mice so as not to artificially bias quantification of overlap, Std pups were given a novel enrichment object (a plastic exercise saucer) in the home cage at P17 prior to 4-OHT administration. We quantified the density of early-life (Std or ELS) activated GFP+ cells, adult experience (control or CNSDS)-activated c-Fos+ cells, and their overlap across key stress-responsive mesocorticolimbic brain regions including NAc, PFC, BLA, and VTA. All statistical analyses including main effects, interactions, and post-hoc analyses are included in **Supplemental Tables 1-2**.

In all brain regions and analyses, there was no main effect of sex and thus male and female samples were analyzed together. In all brain regions, we found similar numbers of GFP+ cells in response to Std and ELS on P17 (**Figures 4A, 4E, 4I, 4M**).

In NAc, we found a main effect of adult stress on cFos+ cells [*F*(1,27)=15.44, *p*=0.0005, **Figure 4B**], density of co-labeled cells [*F*(1,27)=4.581, *p*=0.042, **Figure 4C**], and percent of early-experience-activated cells that were reactivated by adult stress [trend-level: *F*(1,27)=3.610, *p*=0.068, **Figure 4D**]. There were also significant interactions between early-life and adult stress on activation [cFos+ cells: *F*(1,27)=5.339, *p*=0.029], density of co-labeled cells [*F*(1,27)=5.062, *p*=0.033], and percent of early experience-activated cells that were reactivated by adult stress: [*F*(1,27)=4.367, *p*=0.046]). Within ELS-reared mice, but not Std mice, post-hoc analysis with Tukey’s multiple-comparisons correction showed significant effects of adult stress on cFos+ cellular density (*p*<0.0001), density of co-labeled cells (*p*=0.002), and percent of ELS-activated cells reactivated by adult stress (*p*=0.005). Together, these results show a preferential activation (**Figure 4B**) and reactivation (**Figures 4C and 4D**) of ELS-activated cells by adult stress in NAc.

In PFC, we observed that adult stress had a main effect on cFos+ cells [*F*(1,27)=12.28, *p*=0.002, **Figure 4F**], co-labeled cell density [trend-level: *F*(1,27)=0.107, *p*=0.053, **Figure 4G**], and the percentage of early experience-activated cells that were reactivated by adult stress [*F*(1,27)=6.644, *p*=0.016, **Figure 4H**]. There was a trend for a main effect of ELS on PFC ensemble reactivation [*F*(1,27)=3.690, *p*=0.065]. There were also interactions between early-life and adult stress on activation [cFos+ cells: *F*(1,27)=7.261, *p*=0.012] and the percentage of early experience-activated cells that were reactivated by adult stress [*F*(1,27)=5.888, *p*=0.022]. Post-hoc analysis using Tukey’s multiple-comparisons correction revealed significant effects of adult stress on cFos+ cellular density (*p*<0.0001), density of co-labeled cells (*p*=0.009), and percentage of ELS-activated cells reactivated by adult stress (*p*=0.0004) in ELS-reared mice but not Std mice. In PFC, we also found a significant difference in cFos+ cells and percentage of GFP+ cells that were reactivated, between Std and ELS mice who have not experienced adult stress (*p*=0.022 and *p*=0.017, respectively). Together, these findings indicate that adult stress preferentially activates (**Figure 4F**) and reactivates (**Figure 4H**) ELS-activated cells in the PFC. However in PFC, there was only a trending main effect of adult stress on colocalization [trend-level: *F*(1,27)=0.107, *p*=0.053] and no interactions between early-life and adult experience on colocalization, in contrast to NAc.

In BLA, we found strong main effects of adult stress on cFos+ cell density, co-labeled cell density, and the proportion of early experience-activated cells that were reactivated by adult stress [*F*(1,27)=8.441, *p*=0.007, *F*(1,27)=5.42, *p*=0.028, and *F*(1,27)=24.27, *p*<0.0001, respectively, **Figures 4J-L**]. There was a trend-level main effect of ELS on ensemble reactivation in BLA [*F*(1,27)=4.091, *p*=0.053, **Figure 4L**]. Adult stress increased the density of cFos+ cells in ELS mice (*p*=0.0002) as well as in standard-reared mice (trend, *p*=0.051, **Figure 4J**). Adult stress also increased the density of co-labeled cells (*p*=0.017), and the percentage of ELS-activated cells reactivated by adult stress (*p*=0.008) in ELS-reared mice but not in Std mice, as per a post-hoc analysis using Tukey’s multiple-comparisons correction. However, there were no significant interactions between early-life and adult stress within the BLA, indicating that while BLA strong encodes recent adult stress experience, its activity may not encode a memory of prior ELS that contributes to stress sensitization.

Early-life or adult stress had no effect on overall cFos+ activity in VTA (**Figure 4N**) or ELS-activated cell reactivation (**Figures 4O and 4P**). Of the reactivated cells in VTA, we also examined the proportion that were dopaminergic (overlap of TH+, cFos+, and sfGFP+). We found <10% of reactivated cells were dopaminergic, with no significant group differences (data not shown).

### Chemogenetic inactivation of NAc ELS-ensembles rescues behavioral hypersensitivity to adult social stress

Given that ELS-activated cells were preferentially reactivated by adult stress in NAc, we next sought to test whether reactivation of ELS-activated cells in NAc was functionally relevant for hypersensitivity to adult stress at the behavioral level using a chemogenetic approach. NAc also receives glutamatergic inputs from mPFC and BLA where we also found trending main effects of ELS on ensemble reactivation, making it an ideal region to target for functional manipulation. We hypothesized that inhibition of ELS-activated NAc neurons — but not a random subset of NAc neurons — during adult stress would alleviate hypersensitivity to the stressor on behavioral measures.

It was recently shown that olanzapine, a clinically approved antipsychotic drug with a more favorable side-effect profile than clozapine, is a highly potent activator of hM4Di (Lieb et al., 2019; Weston et al., 2019; Upright and Baxter, 2020). OLZ also readily crosses the blood-brain barrier unlike the more common DREADD ligand CNO (Wang et al., 2004; Loryan et al., 2016; Gomez et al., 2017). Since the most common side effects of olanzapine are weight gain and drowsiness (Alper et al., 2007; van der Zwaal et al., 2014), we confirmed there was no effect of OLZ alone on weight or depression- and anxiety-like behaviors prior to using it as a DREADD ligand in our behavioral studies. To examine this, adult male wild-type C57Bl6/J mice were tested for baseline social avoidance and open field exploration, administered i.p. injections of either saline, 0.1 mg/kg OLZ, or 1 mg/kg OLZ 30 minutes prior to CNSDS daily for 10 days, and retested for the same behaviors. Neither dose of OLZ altered weight gain across the 10 days (**Supplemental Figure 1B**), consistent with other mouse work (Shertzer et al., 2010; Weston et al., 2019). Neither the social interaction test (**Supplemental Figures 1C-D**) nor the open-field test revealed any significant differences between mice injected with saline and either dose of OLZ (**Supplemental Figures 1E-F; Supplemental Table 3**). Although it appears that OLZ-treated mice spent less time in the center of an open field than vehicle-treated mice, it is important to note that mice were randomized to treatment without regard for open-field center time, which happened to differ at baseline, and the duration did not differ from the pre-stress test (**Supplemental Figure 1F**). We proceeded with a dose of 0.1 mg/kg OLZ based on previously published hM4di ligand efficacy at this dose (Weston et al., 2019).

To manipulate specific ELS-activated neuronal ensembles hypothesized to contribute to ELS-induced behavioral dysfunction, we injected AAV expressing a Cre-dependent inhibitory DREADD hM4di in NAc of *Arc-CreER^T2/+^* pups at P7 and allowed virus to express for 10 days before inducing recombination in Std or ELS-activated neurons with 4-OHT (**Figure 5 A-C**). We then employed a within-subject test design to assess behavior before adult stress, after CNSDS with vehicle for full experience of social stress, or after a second round of CNSDS with 0.1 mg/kg OLZ (i.p) to inhibit early-life experience-activated ensembles during social stress experience. We found a main effect of sex across behavior tests and analyzed sexes separately. All statistical analyses including main effects, interactions, and post-hoc analyses are included in **Supplemental Tables 4-5**.

There was a significant interaction between ELS and CNSDS with treatment on social avoidance behavior among males [social interaction ratio: *F*(2,22)=4.332, *p*=0.0259; **Figure 5D**]. CNSDS with vehicle treatment decreased the proportion of male mice categorized as resilient among ELS [*z*(1,12)=2.00, *p*=0.045], but not Std males [*z*(1,12)=1.08, *p*=0.280] compared to pre-CNSDS testing levels within each group (**Figure 5E**). DREADD inhibition with OLZ treatment compared to vehicle increased the proportion of ELS mice considered resilient [*z*(1,12)=2.45, *p*=0.014], but not the proportion of resilient Std males (*p*=0.577). These data replicate prior findings (Peña et al., 2017) of an interaction between ELS and adult stress such that two-hits of stress increases social avoidance (**Figure 5F**), and demonstrates efficacy of DREADD inhibition of ELS-activated NAc ensembles for rescuing social avoidance behavior after the second round of CNSDS.

In a novelty-suppressed feeding test, all but one male (Std-OLZ) was observed to eat in the novel arena. Contrary to prior literature, adult stress did not increase latency to eat in the novel arena and there were no main effects of or interactions between ELS and CNSDS with treatment (**Figure 5G**). In an open field test, there was a main effect of CNSDS and treatment on center time [*F*(2,22)=77.48, *p*<0.0001], such that adult stress decreased center exploration. However, there was no interaction with ELS or rescue by DREADD-inhibition of NAc ensembles.

We also examined the behavior of female mice. Female mice spend less time in the center of an open field arena after each round of CNSDS [main effect of CNSDS and treatment: *F*(2,22)=6.388, *p*=0.007], indicating CNSDS was an effective stress in females (**Supplemental Figure 2A**). Post-hoc analysis shows a decrease in open field center time only among ELS females (*p*<0.040), but not Std females (*p*=0.492). However, there were no main effects of, or interactions between, ELS and CNSDS with DREADD inhibition on novelty-suppressed feeding or social avoidance behavior (**Supplemental Figures 2B-D**).

## DISCUSSION

There is a critical gap in our understanding of the mechanisms by which ELS sensitizes response to future stressors. Our objective was to investigate whether this hypersensitivity arose from overactivation of stress-responsive brain regions broadly, or from activity within specific ensembles of ELS-responsive neurons. To overcome the challenge of identifying neurons active during a specific past experience, we leveraged *Arc-CreER^T2/+^* transgenic mice to permanently label early experience-activated neurons and track them across the lifespan. We first showed that distinct neuronal ensembles in NAc respond to distinct early life experiences (**Figure 2C**). We then showed that while adult stress activated neurons in NAc, mPFC, and BLA more strongly than a control experience, these regions were not simply hyperactive in mice with prior ELS experience compared to standard-reared mice (**Figure 4**). Instead, we found that adult stress preferentially reactivated ELS-activated cells in both NAc and mPFC (**Figure 4**), supporting our second hypothesis that specific ensembles of ELS-activated neurons in these regions contribute to lifelong stress hypersensitivity. Finally, inhibition of ELS-activated NAc neurons, but not control-tagged neurons, rescued social avoidance behavior following adult stress in males (**Figure 5**). These data provide evidence that ELS-activated neuronal ensembles in corticolimbic brain regions remain hypersensitive to stress across the lifespan and contribute to behavioral stress sensitivity.

Context and time are both essential factors in the retrieval of cellular memories. Neuronal ensembles active in one context — whether a particular portion of a maze or shock-paired chamber — are less likely to be active in a distinct context (O’Keefe and Nadel, 1978; Josselyn et al., 2015; Moser et al., 2015; Tonegawa et al., 2015a; Cai et al., 2016). Time also erodes memory ensembles, such that there is greater reactivation of hippocampal ensembles in an identical context at 5 days compared to 30 days post-experience (Denny et al., 2014). Memories of experiences close in time are also more likely to share overlapping neuronal ensembles than experiences separated by a week or more (Cai et al., 2016), and longitudinal calcium recordings of cells within prefrontal cortex shows substantial representational drift of stress-sensitive ensembles across days (Patel et al., 2022), which is likely due to destabilization of the ensemble during repeated retrieval (Josselyn et al., 2015; Cho et al., 2021). Given this, the length of time between initial ELS and adult stress exposures, and the different contexts in which each stress was experienced would predict low reactivation of ELS-sensitive cells in the current study. Nevertheless, we found a similar or greater percent reactivation of neurons between ELS and adult CNSDS (~5-15% depending on brain region) as fear conditioning studies find in hippocampus (~5%) and amygdala (~10-15%) (Denny et al., 2014; DeNardo et al., 2019). This enduring encoding of ELS may therefore contribute to stress hypersensitivity and lifetime risk for psychiatric disease.

While we specifically sought to understand how ELS primes cellular activity responses to adult stress, another important question is how stable ensembles of ELS-activated cells are across days of ELS. Earlier work found similar increases in serum corticosterone following a single bout of maternal separation as compared to repeated maternal separation from P1-14 or P14-21, suggesting that pups do not habituate their physiological and hormonal stress response from the first day to the last day of ELS (Horii-Hayashi et al., 2013; Nishi et al., 2013). Horii-Hayashi (2013) previously found that the number of cFos-expressing cells was generally higher following a single bout of maternal separation on P14 or P21 compared to the last day of repeated maternal separation, although significant increases in cFos+ cells were still observed following repeated maternal separation. Our examination of cFos response to ELS on P17 with (chronic) or without (acute) prior ELS showed similar levels of activated cells in both conditions, replicating these earlier findings (**Figure 2B**). Our choice to tag cells on the last day of ELS rather than the first day therefore appeared to capture the same number of ELS-activated cells as were tagged on the first day of ELS, and likely the ones most robustly activated across all days of ELS. Interestingly in adults, mPFC ensembles tagged at the end of fear conditioning are more likely to be reactivated and contribute to behavior, even at remote time points, than those tagged during early stages of learning (DeNardo et al., 2019), further supporting our strategy to induce recombination on the last day of ELS. Finally, we found a higher percentage of cFos+ cells on the last day of ELS that were also active on the first day of ELS, compared to a low percentage of overlap if early experience was mis-matched (**Figure 2C**), indicating that even while the homecage enrichment experience engaged a similar number of neurons across brain regions compared to ELS (Figure 4), these early-life experiences engaged distinct neuronal ensembles.

Heightened neuronal excitability biases recruitment into ensembles (Kida et al., 2002; Han et al., 2008; Sekeres et al., 2010; Kim et al., 2014; Yiu et al., 2014; Josselyn et al., 2015). We did not find evidence that ELS increased numbers of cells expressing cFos at baseline in any brain region (**Figure 4 B, F, J, N**, control), nor reactivation of ELS-ensembles in the absence of adult stress (**Figure 4 C, G, K, O**, control). Interestingly, in some regions cFos density and overlap was lower in ELS-exposed compared to Std-reared mice. One possibility for this effect could be that while Std-tagged cells responsive during positive-valence/novelty experience are more likely to overlap with random homecage activity in adulthood, ELS-tagged cells are not random such that both the lower control levels and the higher CNSDS reactivation indicate specificity. A lack of continual hyperactivity across the lifespan is also consistent with the idea that ELS ensembles are dormant or even suppressed below baseline outside of retrieval by additional stress (Josselyn et al., 2015). Even dormant ensembles have been shown to facilitate response to stimuli through strengthened synaptic connectivity and strength, glutamate receptor expression, expression of the transcription factor CREB, or by enduring epigenetic mechanisms (Kida et al., 2002; Han et al., 2008; Sekeres et al., 2010; Kim et al., 2014; Yiu et al., 2014; Marco et al., 2020). It will be important for future research to investigate which molecular mechanisms ELS-reactivated neurons engage for sensitized stress response.

We found an interaction between ELS and adult stress on neuronal reactivation within both NAc and mPFC, suggesting that multiple brain regions may contribute to hypersensitivity to adult stress. We chose to manipulate ELS-activated cells in NAc, as NAc integrates glutamatergic inputs from PFC and BLA and dopaminergic inputs from VTA as well as hippocampus and other regions (French and Totterdell, 2002, 2003; Goto and Grace, 2008; Britt et al., 2012; Bagot et al., 2015; Ramirez et al., 2015). It is possible that manipulating activity of ELS- and enrichment-activated cells upstream of NAc, in PFC or BLA, would similarly result in behavioral rescue. Indeed, optogenetic inhibition of BLA in juvenile mice exposed to ELS promotes social behavior (Opendak et al., 2021). However, broad inhibition of all glutamatergic activity by optogenetically-induced LTD in mPFC or amygdala projections to NAc prior to behavioral testing did not alter susceptibility or resilience following adult social defeat stress (Bagot et al., 2015).

Chemogenetic inhibition of ELS-activated NAc neurons, but not control-tagged neurons, rescued social avoidance behavior following CNSDS in males (**Figure 5D-F**). To control for effects of vehicle vs ligand within-subject, we employed a repeated testing design wherein mice were exposed to CNSDS twice: first with vehicle administration wherein all mice were assumed to experience the full stress, and a second full 10-days of CNSDS with the DREADD ligand administered daily before each defeat stress exposure. We reasoned that it would be more challenging to rescue behavior after mice had already experienced social defeat and had already displayed behavioral changes associated with susceptibility to stress. While the strongest effect of inhibition was on social avoidance behavior, latency to eat in a novelty-suppressed feeding task appeared to be selectively reduced by DREADD-inhibition in ELS and not Std males. Effective rescue of social avoidance behavior in males and the proportion of mice considered susceptible supports an enduring functional role for ELS-activated NAc neurons in ELS-induced stress hypersensitivity.

While our imaging findings include both males and females, and female ELS-exposed mice appear to respond to CNSDS by reduced open field center time, there was no effect of NAc ensemble inhibition on female behavior across three tests. Recent work using machine learning-based behavioral analysis of female behavior during social defeat attacks also shows relatively more subtle behavioral responses to defeat compared to males (Willmore et al., 2022). Additional research is needed to determine whether distinct adult stressors would produce a more robust effect, and whether other brain regions not assessed in the current study such as within the hypothalamus more robustly encode female cellular memory of ELS.

Heterogeneity of neuronal subtype in NAc and other brain regions is important to consider, as Drd1- and Drd2-expressing medium spiny neurons (D1- and D2-MSNs) have been found to contribute to different aspects of reward and aversion, with D1-MSNs classically thought to encode reward signals and D2-MSNs encoding aversive signals (Hikida et al., 2010; Kravitz et al., 2012; Kupchik et al., 2015; Kupchik and Kalivas, 2017; Soares-Cunha et al., 2020). While we took an activity-dependent rather than a molecularly-defined approach and did not distinguish between D1- and D2-MSNs in NAc in the current study, recent work found that ELS predominately impacts the molecular profiles of D2-MSNs in NAc (Kronman et al., 2021). Consistent with a hypothesis that ELS predominantly acts on D2-MSNS, adult stress in male mice has been found to increase excitatory transmission onto D2-MSNs and decrease excitatory transmission onto D1-MSNs (Francis et al., 2015). While inhibiting D1-MSNs in stress-resilient mice increases depression-like behavior, and activating D2-MSNs prior to stress increases susceptibility, the effect of inhibiting D2-MSNs specifically during stress experience has not yet been reported and would be an important follow-up for the current study (Francis et al., 2015; Peña, 2017).

In sum, these data provide evidence that ELS-induced stress hypersensitivity is encoded at the level of neuronal ensembles in NAc. This work paves the way for additional research to understand how activation of corticolimbic neurons by ELS alters molecular development and circuit integration of these cells, and to formulate translational strategies to ameliorate stress hypersensitivity in those who experienced ELS and are at risk for developing mood and anxiety disorders.

## ACKNOWLEDGEMENTS

This research was funded by NIH R00MH115096 (CJP); NIH R01MH129643 (CJP); PNI Research Innovator Award (CJP); NYSCF (CJP). CJP is a New York Stem Cell Foundation Robertson Investigator.

We would like to thank Dr. Abigail Polter for helpful early discussion, and Tanzina Islam and Dana Waitman for their assistance with images. Parts of some figures were generated with BioRender.com.

## AUTHOR CONTRIBUTIONS

CJP and JAB designed the studies. JAB, CM, AM, and RR collected the data. SB maintained the transgenic lines and genotyped mice. JAB, CM, and CJP analyzed the data and created figures. JAB and CJP wrote the manuscript with input from all authors. All authors approve the manuscript.

---

**Supplemental Figure 1.**
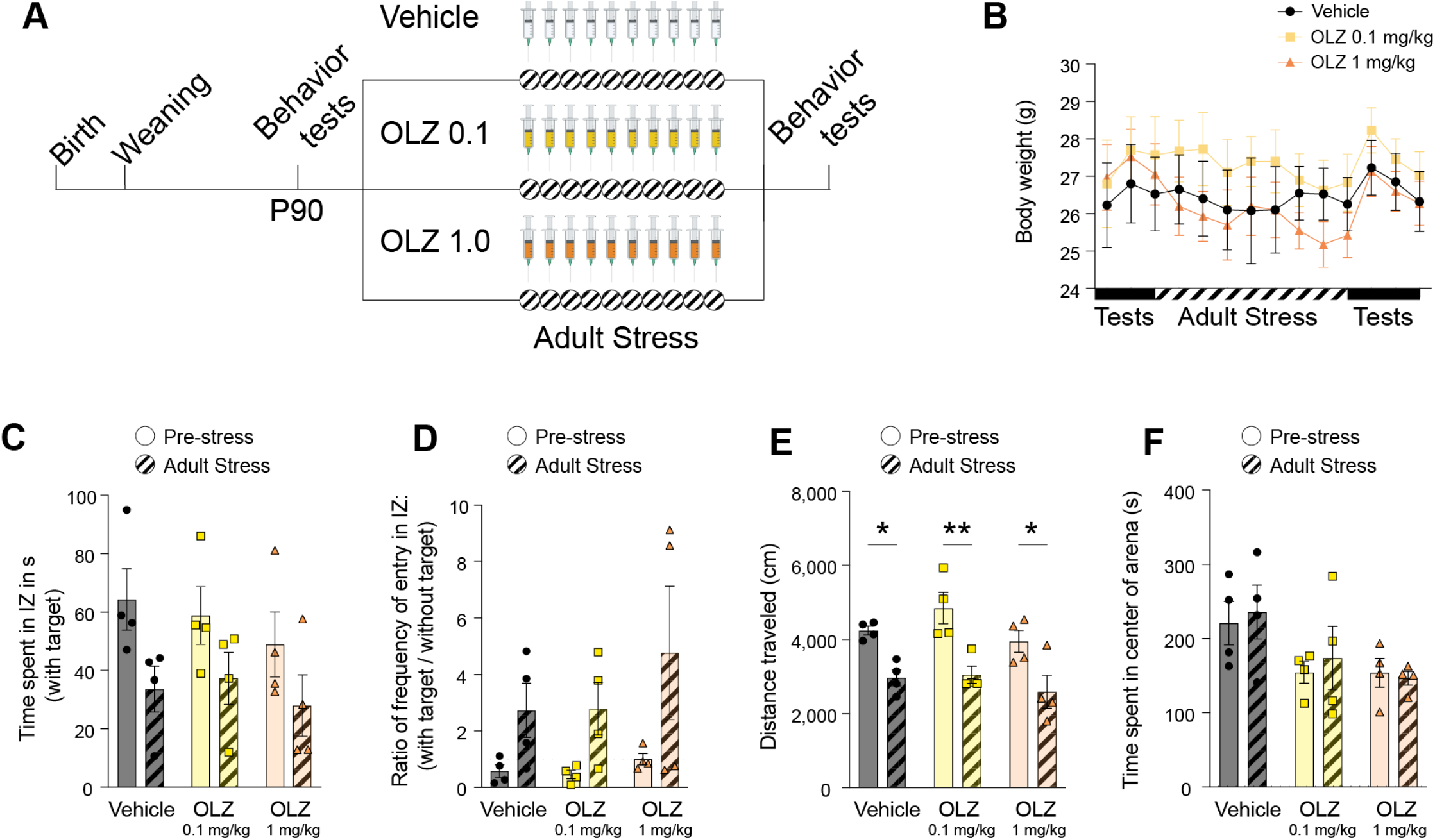
No significant effects of olanzapine treatment by itself on behavior at two doses. (A) Schematic of experimental groups: for 10 days, adult wild-type mice underwent male-only CSDS daily, 30 minutes after receiving an intraperitoneal injection of saline, OLZ (0.1 mg/kg) or OLZ (1 mg/kg). (B) No weight change was observed after 10 days of OLZ injection at either dose selected. In the social interaction test, there were no significant differences between treatment groups in either the time spent in interaction zone (C), or the frequency of entry in the interaction zone (D). In the open field test, all the mice traveled significantly less after defeat (E) but there were no significant differences in time spent in the arena center (F).

**Supplemental Figure 2.**
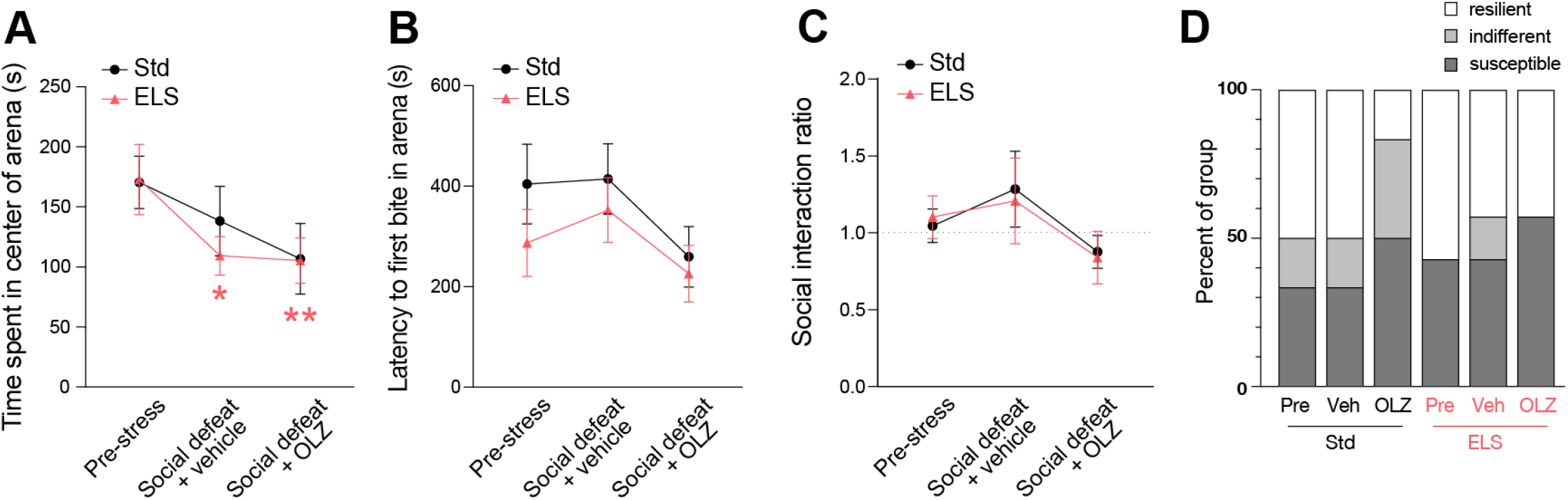
Female behavior remained largely unchanged by CNSDS and DREADD inhibition of NAc ensembles. (A) In the open field test, there was a main effect of CNSDS and treatment on open field center time among female mice [*F*(2,22)=6.388, *p*=0.007]. However, there was no interaction with ELS and no rescue of behavior by OLZ treatment in post-hoc comparisons. Post-hoc comparisons between corresponding conditions and pre-stress data are indicated by *(*p*<0.05) and **(*p*<0.01). In the novelty-suppressed feeding (B) and the social interaction test (C-D), no significant differences were observed between ELS and standard-reared mice.

**Supplemental Table 1**: Statistical analysis related to Figure 4 GFP comparisons.

**Supplemental Table 2**: Statistical analysis related to Figure 4 ANOVAs.

**Supplemental Table 3**: Statistical analysis related to Supplemental Figure 1 OLZ pilot.

**Supplemental Table 4**: Statistical analysis related to Figure 5 male behavior.

**Supplemental Table 5**: Statistical analysis related to Supplemental Figure 2 female behavior.

